# Differential Susceptibility of Rat Retinal Ganglion Cells Following Optic Nerve Crush

**DOI:** 10.1101/429282

**Authors:** Kirstin B. VanderWall, Bin Lu, Shaomei Wang, Jason S. Meyer

## Abstract

Retinal ganglion cells (RGCs) are a heterogeneous group of cells, comprised of numerous subpopulations, that work together to send visual information to the brain. In numerous blinding disorders termed optic neuropathies, RGCs are the main cell type affected leading to degeneration of these cells and eventual loss of vision. Previous studies have identified and characterized RGC subtypes in numerous animal systems, with only a handful of studies demonstrating their differential loss in response to disease and injury. Thus, efforts of the current study utilized an optic nerve crush (ONC) model to characterize the loss of RGCs and disease phenotypes associated with this injury. Additionally, the loss of RGC subtypes including direction selective-, alpha-, and ip-RGCs following ONC was explored. Results of this study demonstrated the differential loss of RGC subtypes with a high susceptibility for loss of alpha- and direction selective-RGCs and the preferential survival of ip-RGCs following ONC and allows for the establishment of additional studies focused on mechanisms and loss of these cells in optic neuropathies. Additionally, these results put important emphasis on the development of therapeutics targeted at the loss of specific subtypes as well as cellular replacement following injury and disease.

## Introduction

Retinal ganglion cells (RGCs) are the primary projection neuron of the retina and extend axons through the optic nerve to synapse with post-synaptic targets in the brain (Levin, 2005). A disruption in this crucial connection due to injury or optic neuropathies can result in the degeneration of RGCs and loss of vision or blindness (Agostinone and Di Polo, 2015). Although most RGCs serve a similar function, to transmit visual information to the brain, this class of cells has been proven to be highly diverse, with the classification of more than 30 subtypes to date (Berson, 2008; Dhande et al., 2015; Sanes and Masland, 2015). RGC subtypes have been previously classified based on differences in morphological features and functional capabilities, collectively contributing to the multifaceted integration of visual stimuli. The identification of RGC subtypes has been achieved through the expression of specific molecular signatures (Boycott and Wassle, 1974; De la Huerta et al., 2012; Diao et al., 2004; Fukuda, 1977; Trimarchi et al., 2007) in a variety of animal models including mouse, rat, and cats with a small number of studies identifying RGC subtypes in human derived cells (Daniszewski et al., 2018; Langer et al., 2018; Ohlemacher et al., 2016; Sluch et al., 2015). The ability to successfully identify RGC subtypes allows for more in-depth studies of their development and contribution to vision. Furthermore, an important precedence stands to recognize how various RGC subtypes exhibit differential survival following disease or injury in order to better understand disease mechanisms and develop more targeted treatments.

More recently, studies have demonstrated the identification and subsequent degeneration and death of various RGC subtypes in various injury and disease models, including axotomy, optic nerve crush, and glaucoma models (Birke et al., 2010; Cui et al., 2015; Daniel et al., 2018; Duan et al., 2015; El-Danaf and Huberman, 2015; Majander et al., 2017; Ou et al., 2016; Puyang et al., 2017; Yucel et al., 2003), suggesting subtype specific responses to injury. However, these few studies produced limited conclusions as to how various model systems affect RGCs and more specifically, which RGCs subtypes are highly affected in response to disease and injury. Additional studies depicting the precise loss of RGC subtypes and the extent of this loss would provide necessary insight into the development of specific therapeutics, with important implications for targeted RGC subtype replacement for injury and optic neuropathies in future studies.

Thus, initial efforts of this study focused on the examination of RGC loss and disease phenotypes associated with optic nerve crush (ONC) injury which may elucidate similar features to those observed during glaucomatous neurodegeneration. Initially, rat retinas were characterized for various disease phenotypes following injury. ONC resulted in a significant loss of RGCs and induced disease phenotypes including glial reactivity, increased apoptosis, and decreased behavioral response. Additionally, specific RGC subtypes were examined after ONC and demonstrated a significant loss in different classes of RGCs. Results of the study suggest a high susceptibility for both direction selective-RGCs and alpha-RGCs, and suggest a resilience for non-image forming intrinsically photosensitive-RGCs (ip-RGCs) following ONC. The results of this study provide additional insights into how injury and disease may affect the differential survival of RGC subtypes and the disease phenotypes. More so, these results will aid in the development of therapeutics targeted at specific subtypes, with important implications for future cell replacement therapies of specified subtypes lost in optic neuropathies.

## Results

### ONC results in RGC loss and disease phenotypes

RGCs form the crucial connection between the eye and the brain, allowing for the ability to see (Levin, 2005). In optic neuropathies, RGCs are the main cell type affected with this essential connection severed, leading to profound RGC degeneration and eventual loss of vision (Agostinone and Di Polo, 2015). As the mechanisms and patterns of RGC loss remain inconclusive, further studies are needed for the development of new therapeutics for treatments of optic neuropathies. Efforts of the current study focused on the identification of RGC subtypes in a rat model and characterizing disease deficits following ONC, with important emphasis on the degeneration of RGC subtypes.

Rat retinas were investigated for loss of RGCs at 7 days-post ONC. Immunocytochemistry using RGC-associated protein, RBPMS, demonstrated a significant decrease in the average number of RGCs within the ganglion cell layer following ONC (Figure 1 A-C). ONC animals also displayed lower optokinetic response (OKR) compared to wildtype animals, indicative of some behavioral deficits after injury (Figure 1 D). In addition to RBPMS quantification using ICC, stereology was used as an alternative, unbiased approach to counting RGCs using numerous 100μm squares throughout retinal whole mounts (Figure 1 E). ONC retinas displayed a notable decrease in the number of RGCs per each 100μm square 7 days post injury (Figure 1F). These results indicated a successful injury model using ONC, which resulted in both a significant loss of RGCs with the demonstration of behavioral deficits.

**Figure 1:**
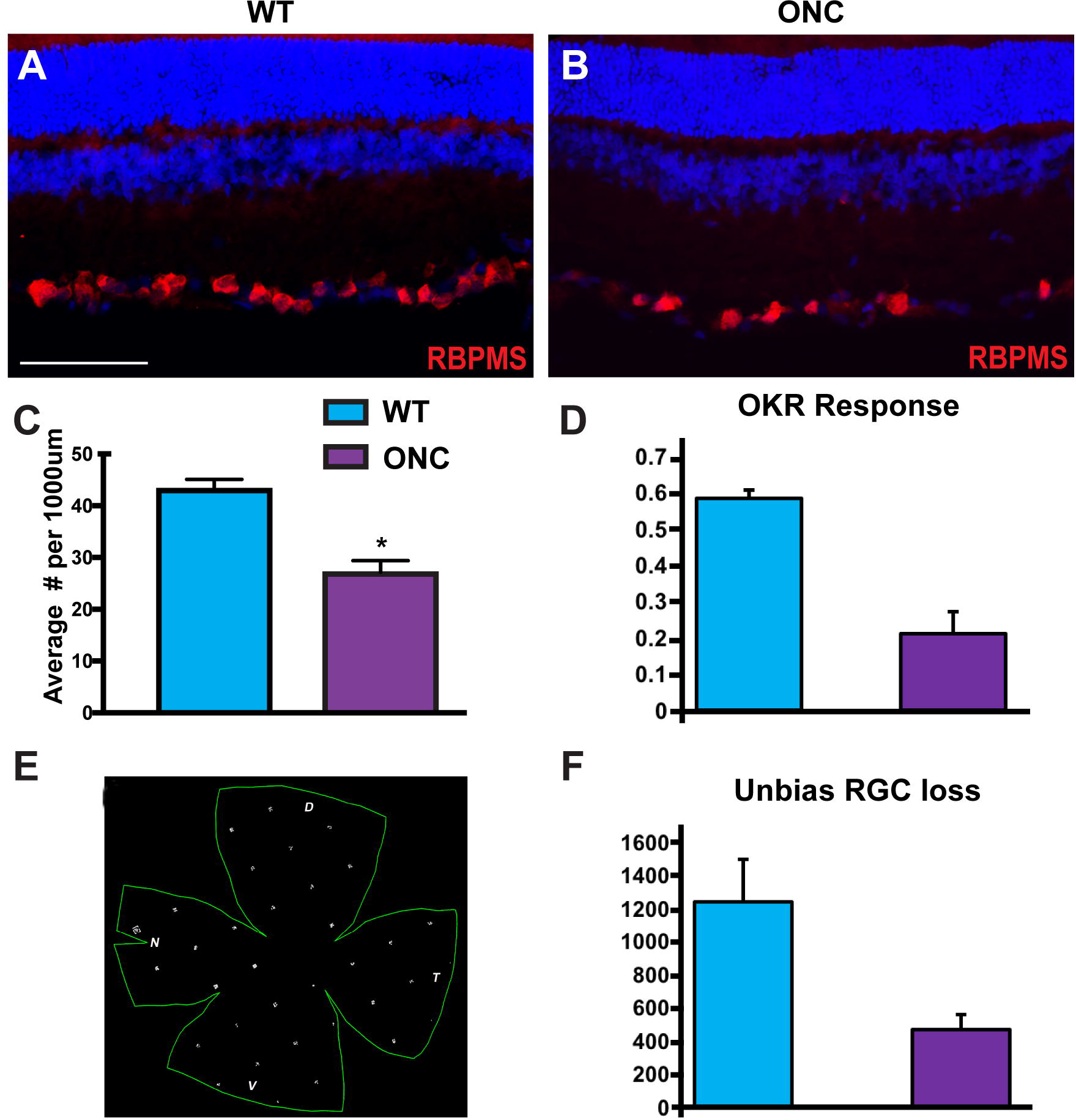
ONC results in profound RGC loss and behavioral deficits. (A). Rat retinal sections expressed RBPMS within innermost layers. (B-C) The average number of RBPMS significantly decreased 7 days after ONC. (D) Behavioral experiments showed decreased optokinetic response (OKR) of animals exposed to injury. (E) Schematic depicting stereology used as an unbiased approach to count RGCs. (F) RGCs counted using stereology were significantly decreased in ONC animals compared to WT animals. Error bars represent S.E.M. Scale bars equal 100μm. Statistical differences were determined using 95% confidence.

Furthermore, rat retinas 7 days-post ONC were analyzed for various other phenotypes typically associated with injury and disease (Libby et al., 2005; Mac Nair and Nickells, 2015; Seitz et al., 2013; Yuan and Neufeld, 2001). One characteristic that has been observed within the CNS following disease and injury is glial activation (Hernandez, 2000; Prasanna et al., 2011; Tezel et al., 2001; Wang et al., 2000). This gliosis normally functions as a safety mechanism in response to an assault, and attempts to halt any further damage to the affected area, as well as provide support to the area in order to heal damage that has occurred. Following ONC, rat retinas exhibited upregulation of GFAP (Figure 2 A-B) within the nerve fiber layer, a characteristic indicative of astrocyte reactivity. Additionally, compared to the wildtype, ONC animals demonstrated microglia infiltration and morphological changes in layers closest to the ganglion cell layer, characteristics related to microglia activation (Figure 2 C-D). Apoptotic activation was examined by the expression of cleaved caspase-3 and was notably higher in ONC RGCs than wildtype RGCs indicating increased cell death within the inner most layers (Figure 2 E-F). RGC loss and ganglion cell layer thinning are also a common characteristic observed in the disease progression of optic neuropathies as well as following injury. In the ONC animals, the average number of nuclei within the ganglion cell layer was significantly lower displaying a thinning of this layer in addition to the downregulation of RGC-associated proteins like RBPMS (Figure 2 G). Thus, these results demonstrate a variety of disease phenotypes associated with ONC injury after 7 days and allow for the further investigation into specific RGC loss in this model system.

**Figure 2:**
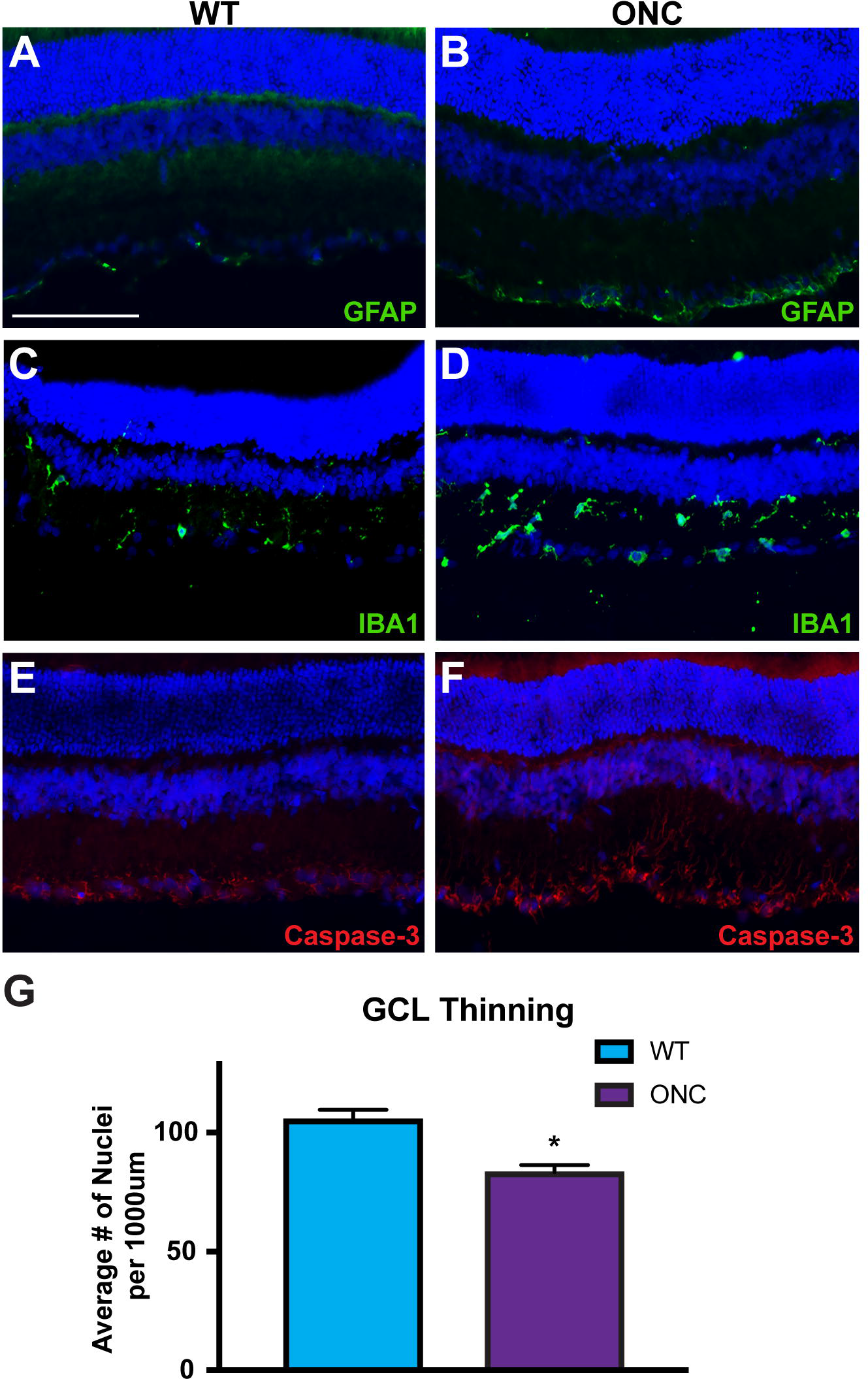
ONC leads to numerous disease-related phenotypes. (A-B) Expression of GFAP was displayed in close association with RGCs in the ganglion cell layer and was notably higher following ONC. (C-D) Microglia were identified near RGCs through the expression of IBA1, with an increased number of microglia as well as a change in morphology found in ONC animals. (E-F) Increased apoptosis was observed within the ganglion cell layer after ONC compared to the WT by the expression of cleaved caspase-3. (G) ONC resulted in significant thinning in the ganglion cell layer demonstrated by decreased average counts for nuclei within inner most layers. Error bars represent S.E.M. Scale bars equal 100μm. Statistical differences were determined using 95% confidence.

### Rat retinas exhibit differential loss of RGC subtypes following ONC

RGCs are a highly diverse population of cells which serve the common purpose to transmit visual information to the brain. Up until recently, the separation of RGCs into specific subtypes was achieved by morphological and functional studies, with each type of RGC exhibiting various dendritic shapes, sizes, and stratification as well as different physiological response to visual information. The heterogeneous nature of RGCs has become increasingly more apparent in recent years with the discovery of novel molecular markers which identify specific populations of subtypes. RGC subtypes have been classified into a few major groups which can be further divided into more than 30 subtypes based on the combinatorial expression of different molecular markers (Dhande et al., 2015; Sanes and Masland, 2015).

The first RGC subtype explored were the alpha-RGCs, which are a group of cells that have various responses to light and dark stimuli and can be broken down into 3 subsets; ON, Sustained OFF, and Transient OFF based on the combinatorial expression of different molecular markers (Jelinek et al., 2011; Krieger et al., 2017; Pang et al., 2003; Peichl, 1991). Osteopontin and SMI32 have been established as pan-identifiers of alpha-RGCs, and as such SMI32 was utilized to visualize and quantify this class of cells. ICC of WT and ONC retinas demonstrated expression of SMI32 selectively within inner most layers (Figure 3 A). Quantification of SMI32 7 days-post ONC revealed a significant decrease in the average number of alpha-RGCs within the ganglion cell layer (Figure 3 B-C). These results suggest a heightened susceptibility for the loss of these cells following insult and disease.

**Figure 3:**
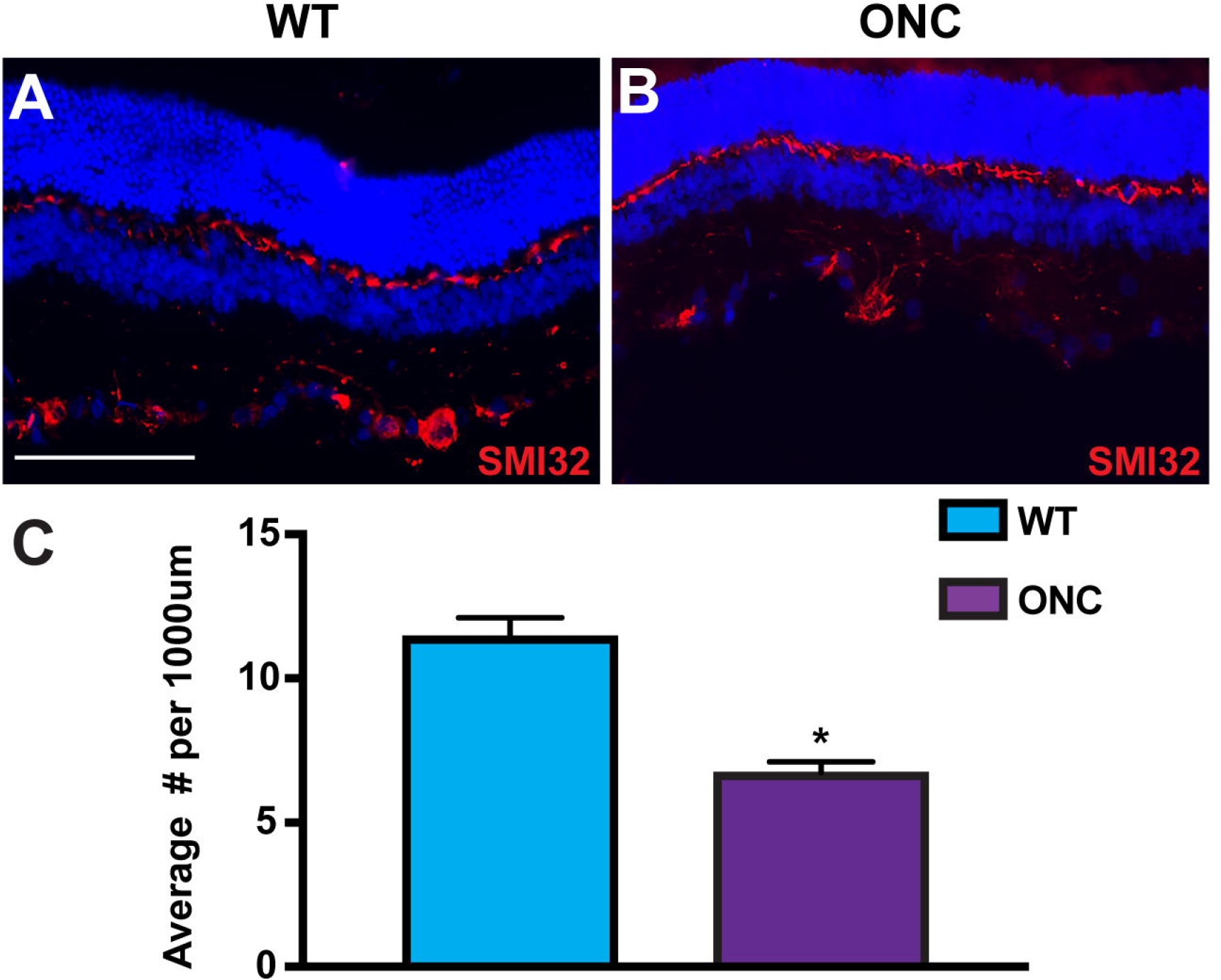
Alpha-RGCs display susceptibility to injury. (A) Alpha-RGCs were identified within the ganglion cell layer by the expression of pan-marker SMI32. (B-C) Following ONC, the expression of SMI32 was significantly decreased compared to WT animals. Error bars represent S.E.M. Scale bars equal 100μm. Statistical differences were determined using 95% confidence.

Direction selective-RGCs were also investigated within rat retinas. These cells have the unique ability to respond to preferred directional motion in response to light and dark stimuli (Cruz-Martin et al., 2014; Liu, 1995). Numerous molecular markers have been established to identify these cells, with the combinatorial expression of numerous markers used to characterize specific subsets of direction selective-RGCs (De la Huerta et al., 2012; Duan et al., 2014; Huberman et al., 2009; Sanes and Masland, 2015). Two types of direction selective-RGCs were investigated in this study, ON-OFF direction selective-RGCs identified by CART, and a subset of ON direction selective-RGCs marked by FSTL4. CART and FSTL4 were highly expressed within the ganglion cell layer of rat retinas (Figure 4 A-B). When analyzed 7 days-post ONC, both types of direction selective-RGCs were significantly decreased within the inner most layers compared to WT animals (Figure 4 C-E). Thus, these results revealed a high degree of susceptibility for direction selective-RGCs following injury.

**Figure 4:**
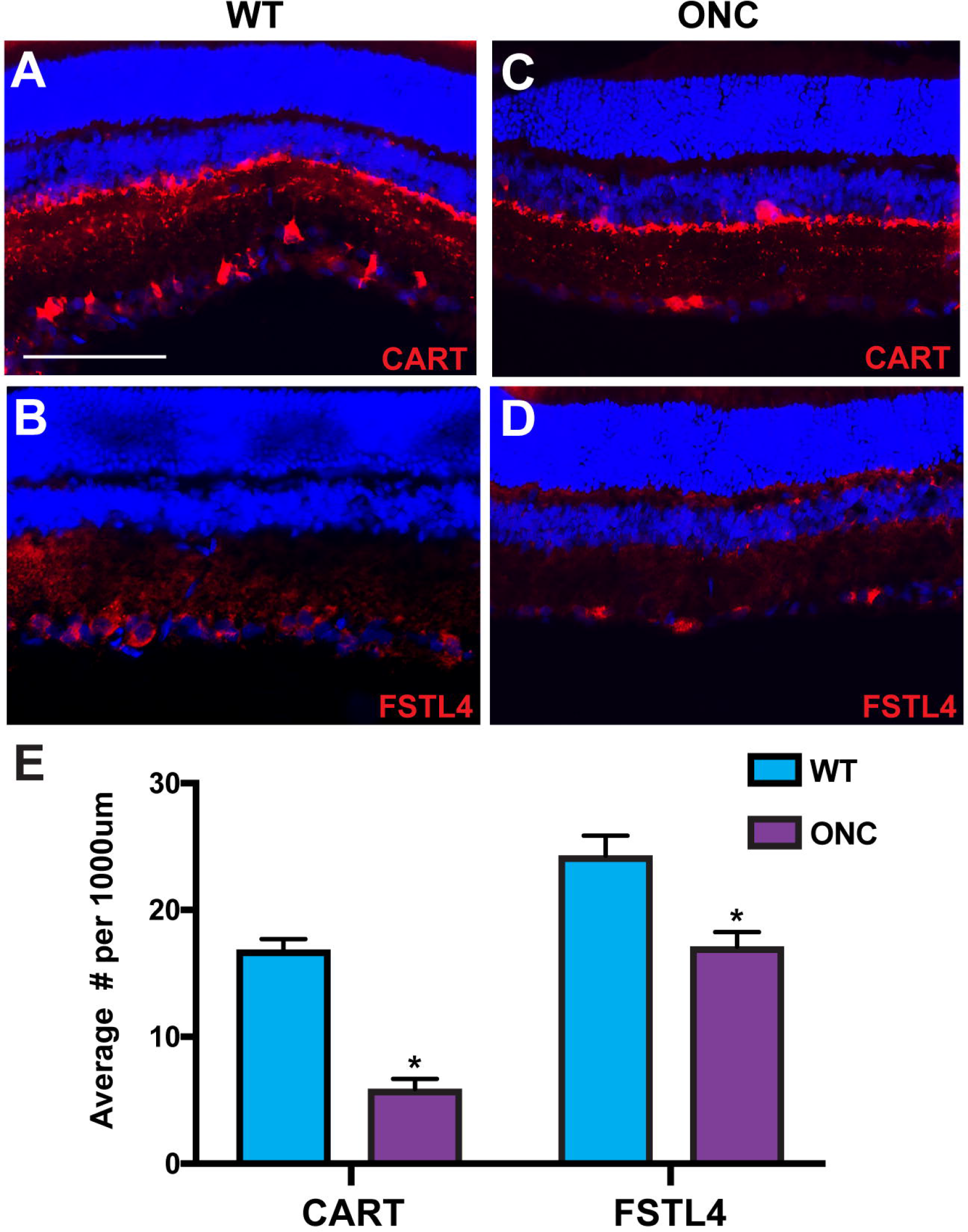
ONC results in a severe loss of direction selective-RGCs. (A) CART expression was displayed and identified ON-OFF direction selective-RGCs in rat retinas. (B) A subset of ON direction selective-RGCs were marked by FSTL4 and indicated high levels of expression within the ganglion cell layer. (C-E) Following ONC, both CART and FSTL4 demonstrated a significant reduction in their average number per retina length. Error bars represent S.E.M. Scale bars equal 100μm. Statistical differences were determined using 95% confidence.

Lastly, the loss of ip-RGCs was explored using the ONC model. This class of RGCs demonstrates the distinct ability to respond directly to light stimulus, which aids in the formation of circadian rhythms and pupillary reflexes. Previous studies have demonstrated the presence of multiple ip-RGC subtypes which vary in molecular nature and functional capabilities (Chew et al., 2017; Graham and Wong, 1995; Hannibal et al., 2017; Kelbsch et al., 2016; Kofuji et al., 2016; Sweeney et al., 2014). In this study, ip-RGCs were readily identified by the expression of TBR2 and Melanopsin (OPN4) within the ganglion cell layer (Figure 5 A-B). Following ONC, no significant change was detected in the average number of TBR2 after 7 days-post ONC (Figure 5 C, E). More so, OPN4 demonstrated a similar level of expression within the ganglion cell layer after 7 days. These results suggest the ability for ip-RGCs to exhibit preferential survival following injury. Taken together, experiments conducted in the current study demonstrated the presence of numerous RGC phenotypes associated with ONC, with an important emphasis on the differential loss and susceptibility of major RGC subtypes.

**Figure 5:**
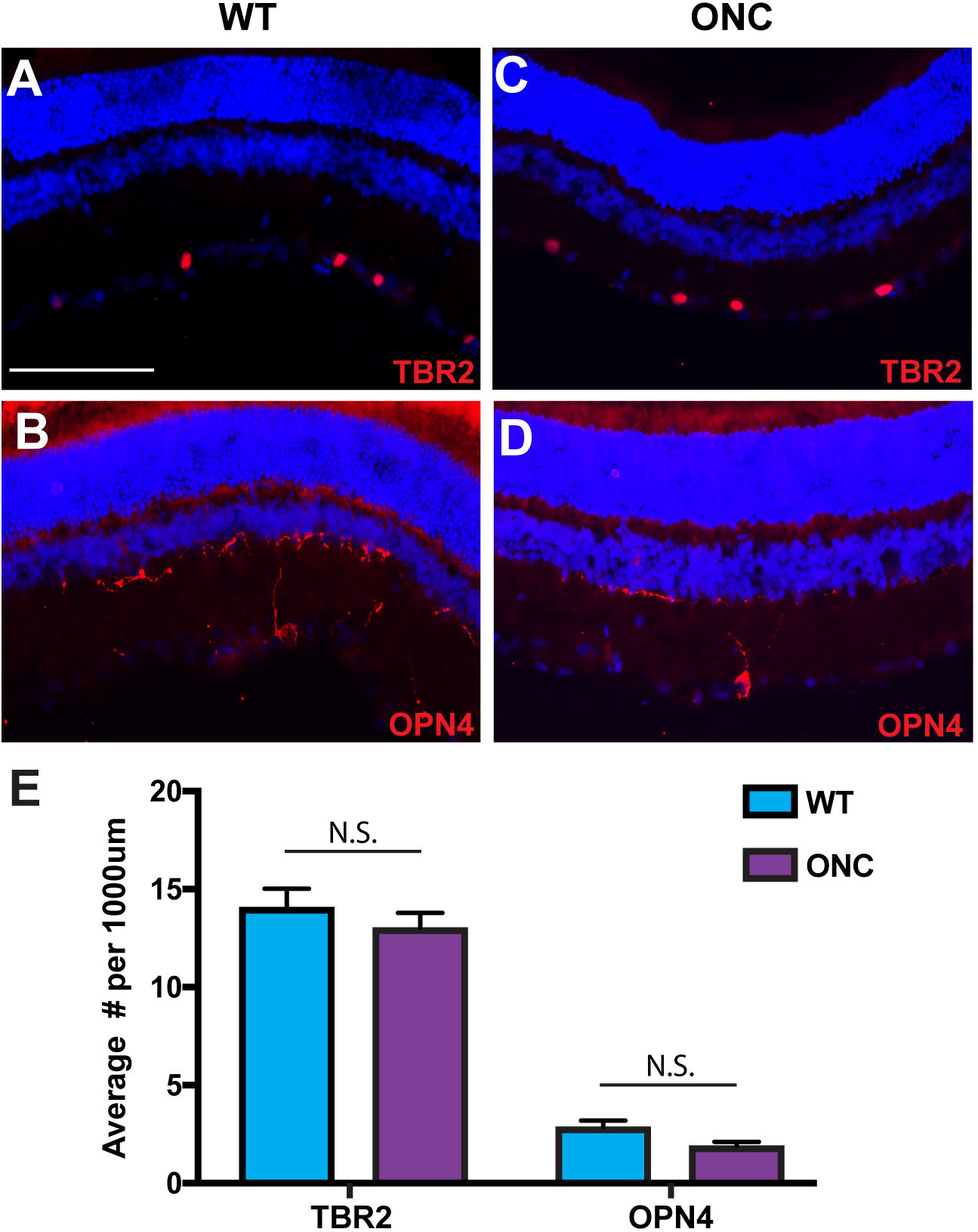
ip-RGCs demonstrate differential survival in response to ONC. (A-B) ip-RGCs were identified in rat retinas by the expression of TBR2 and OPN4. (C-E) TBR2 and OPN4 both retained a similar levels of expression 7 days-post ONC. Error bars represent S.E.M. Scale bars equal 100μm. Statistical differences were determined using 95% confidence.

## Discussion

RGCs are the main cell type affected in optic neuropathies, with degeneration of these cells leading to profound phenotypes and eventual loss of vision. Such diseases are rapidly increasing in their prevalence in the United States with a projected 111.8 million people affected by 2040 (Tham et al., 2014). Therefore, a need exists to develop new treatments and therapeutics targeted at specific disease mechanisms causing the degeneration of RGCs. Previous studies have elucidated the diverse nature of RGCs, with numerous different subtypes discovered to date with differing morphological, functional, and molecular features (Dhande et al., 2015; Sanes and Masland, 2015). More so, recent studies have also suggested a differential susceptibility of RGC subtypes in disease or injury states (Birke et al., 2010; Cui et al., 2015; Daniel et al., 2018; Duan et al., 2015; El-Danaf and Huberman, 2015; Majander et al., 2017; Ou et al., 2016; Puyang et al., 2017; Yucel et al., 2003), providing a new opportunity of targeting specific subtypes for treatments and therapeutics. Efforts of the current study utilized an ONC model in rats to elucidate the degeneration of RGCs following injury. Results demonstrated a significant loss of RGCs as well as their behavioral characteristics following injury. More so, the investigation of major RGC subtypes following injury indicated a heightened susceptibility of direction selective-RGCs and alpha-RGCs, while ip-RGCs displayed a preferential survival compared to those other subtypes. As such, results of this study provide further insight into which subtypes may be most affected in disease and injury states, contributing to the development of treatments and future cell replacement strategies targeted at specific subtypes

The most widely studied phenotype associated with disease and injury to the retina is profound degeneration and loss of RGCs (Agostinone and Di Polo, 2015; Rokicki et al., 2007). Rat retinas, in this study, were investigated for their loss of RGCs following ONC and displayed a significant loss of RGCs in both immunocytochemistry and stereology quantification. More so, RGCs demonstrated decreased behavioral characteristics following ONC with animals displaying lower optokinetic responses, which indicated that RGCs are degenerating as well as losing some of their functional capacities. More so, ONC resulted in presumptive injury responses including reactive gliosis, microglia activation and infiltration, and increased apoptosis, all of which are characteristics associated with disease progression observed in optic neuropathies (Mac Nair and Nickells, 2015; Prasanna et al., 2011; Seitz et al., 2013). These results demonstrated the effectiveness for ONC to serve as an effective injury model and displayed numerous phenotypes known to be associated with disease progression of glaucoma and other optic neuropathies.

Previous studies have used various injury and disease models in different animals to elucidate the loss of specific RGCs subtypes, with results somewhat variable based on the model system or the organism used (Birke et al., 2010; Cui et al., 2015; Daniel et al., 2018; Duan et al., 2015; El-Danaf and Huberman, 2015; Majander et al., 2017; Ou et al., 2016; Puyang et al., 2017; Yucel et al., 2003). Results of the current study demonstrated a high susceptibility for direction selective-RGCs and alpha-RGCs following injury, with their survival significantly decreasing after 7 days post ONC. This phenomenon has been observed in a number of other studies investigating direction selective-RGCs and alpha-RGCs following disease and injury, although some of these studies were looking at specific subsets of each of these major categories and not their loss as a whole. The loss of these two subtypes in our study validates their susceptibility to disease and sets the precedent for futures studies understanding disease mechanisms causing their degeneration, as well as studies targeted at direction selective-RGCs and alpha-RGCs for therapeutic intervention.

In contrast to the high susceptibility of certain RGC subtypes, non-image forming ip-RGCs exhibited differential survival following injury compared to image forming direction selective- and alpha-RGCs in the current study. The preferential survival of ip-RGCs has been observed in other studies which utilized rat and mice injury and disease models, with this subtype in particular demonstrating resilience to insult (Cui et al., 2015; Daniel et al., 2018; Li et al., 2006; Li et al., 2008; Perez de Sevilla Muller et al., 2014; Wang et al., 2018). ip-RGCs are unique in that they have the ability to respond to light using the photopigment, Melanopsin, and play a role in non-image forming functions including circadian rhythms and pupillary reflexes by projecting to the superior colliculus, a non-image forming brain region (Chew et al., 2017; Hannibal et al., 2017; Kelbsch et al., 2016). Due to the unique nature of their functionality, this may lead to their resistance to insult as they aren’t necessary for image formation during visual transduction.

In the current study, ONC was implemented as a model of RGC loss and degeneration. Although this technique is commonly used to model mechanisms resulting in the death of RGCs, many other models exist that can model RGC loss in disease and injury states. The bead occlusion model of glaucoma is a technique used to mimic the gradual increase in intraocular pressure observed in forms of glaucoma. This model differs from the ONC model as it provides a realistic representation of glaucomatous disease manifestation through the progressive increase in pressure resulting in the degeneration of RGCs over time. In future studies, this method should be considered for the study of RGC subtype loss in glaucoma due to its translational similarity to the clinical features observed in various forms of glaucoma.

Optic neuropathies are a class of neurodegenerative diseases that result in the gradual loss of RGCs and vision. As these diseases have a sizeable impact on the quality of life, a need stands for the development of targeted therapeutics to aid in patients experiencing such diseases. As this study, and other studies, have shown, RGC subtypes exhibit differential survival in response to disease and injury, leaving the opportunity to develop more targeted treatments for disease phenotypes. Human pluripotent stem cells (hPSCs) provide an advantageous model for studying RGC subtypes in healthy and disease states as they have been shown to differentiate into a number of these major groups. Furthermore, hPSCs can be used to model optic neuropathies when derived from patients harboring disease causing mutations, allowing for the ability to study the development and degeneration of these subtypes in a human model. In future studies, RGC subtypes differentiated from hPSCs could be used for cellular replacement in late stage disease progression of optic neuropathies, with the potential to purify certain subtypes that may be most susceptible to death and replace those cells. Transplantation of hPSC-derived RGC subtypes into animal models of retinal disease and injury will provide the essential foundation for the development of these strategies for human disease.

Efforts of this study were focused on characterizing the loss of RGCs in an ONC model, with an emphasis on the differential survival of RGC subtypes following injury. Results demonstrated a significant loss of RGCs with the heightened susceptibility of direction selective- and alpha-RGCs following injury with these subtypes significantly decreased by 7 days post ONC. Furthermore, ip-RGCs displayed the ability to survive within the rat retina following injury, indicating a certain resistance in response to disease and injury. Results of the current study provide confirmatory data signifying that RGC subtypes exhibit differential survival following injury and further deliver crucial information in understanding disease mechanisms of RGC degeneration in disease and injury. More so, this study provides further foundation for the development of future therapeutics targeted at the loss of specific RGC subtypes in optic neuropathies.

## Methods

### Animal Care/Optic Nerve Crush Procedure

The Long Evans rats were anesthetized via IP injection of anesthetic agents, followed by carprofen via SQ. Only one eye was operated, the fellow eye was used as untreated controls. Animal's head was fixed in a headholder and their intraorbital optic nerves were exposed after a lateral canthotomy and limbal peritomy. The optic nerve was pinched 1 mm behind the globe for 5 seconds with a special forceps. Special care was taken not to damage the blood supply to the eye traveling along the inferior side of the optic nerve. The retina was examined immediately by ophthalmoscope to ensure blood patency. The right optic nerve (ON) was surgically exposed in its intraorbital segment, which spans 2 to 3 mm beyond the eye cup, taking care to leave the retinal vascularization intact. The lacrimal gland will be displaced, but left intact, and the superior extraocular muscles will be spread to allow access to the ON. An incision was made in the eye-retractor muscle and the meninges perpendicular to the axonal orientation, extending over one third of the dorsal ON. The ON was pinched with a special forceps at a distance of 1 mm behind the eye, leaving the retinal vascularization unaffected. All microsurgery was performed with the aid of a surgical microscope. After all procedures was finished, a reversal agent was given via IP, ophthaine and vetropolycin were applied topically, rats were placed under a warm lamp and monitored until they have recovered from anesthesia.

### Immunocytochemistry

Rat retinal sections were permebalized with 0.2% Triton-X-100 for 10 minutes at room temperature, followed by a wash with 1x PBS. Slides were then blocked using 10% Donkey Serum for 1 hour at room temperature. Primary antibodies were prepared in 0.1% Triton-X-100 and 5% Donkey Serum and applied to the slides overnight at 4°C (RBPMS Phospho Solutions 1830-RBPMS 1:500, CART Phoenix Pharm H-003-62 1:1000, SMI-32 Calbiochem Millipore NE1023 1:500, TBR2 Abcam ab23345 1:500, Melanopsin ThermoScientific PA5-34974 1:500, IBA1 Abcam ab5076 1:200, GFAP Millipore MAB360 1:200, FSTL4 Novus 91913 1:500). The following day, the primary antibodies were removed from the slides with 3 washes of 1x PBS. Then samples were blocked with 10% Donkey Serum for 10 minutes at room temperature. Secondary antibodies were prepared in 0.1% Triton-X-100 and 5% Donkey Serum and applied the slides for 1 hour at room temperature in the dark. The secondary was removed by 3 washes of 1x PBS and slides were properly mounted using appropriate cover glass and poly-mount mounting medium.

### Immunocytochemistry Quantification and Statistical Analyses

RGC-associated and RGC subtype-specific antibodies were quantified from n>2 biological replicates and n=4 technical replicates per animal. Each marker was counted using ImageJ cell counter plugin and the retina length was calculated using appropriated scaling in Image J. Statistical differences were calculated using GraphPad Prism software by a student’s t-test, excluding outliers, and using a 95% confidence.

### Stereology

Stereology (Optical Fractionator) was used as described previously (20) to count the number of RGC in all conditions to minimize bias and optimize reliability. Using an Axioskop 2 MOT Zeiss microscope with a x2.5 objective lens (Zeiss, Germany), a semiautomatic stereology system (StereoInvestigator, Microbrightfield) was used to trace the area of interest, and a x40 magnification (Leitz) was used to count the number of RGC. For each section the computer randomly placed a 100 μm by 100 μm counting frame in different areas of the whole retina (1200 μm X 1200 μm in “Fractioner”) with a number of average sampling sites of 30-40. Cells that were within the counting frame or touching the green line were counted and marked with a cross. The optical fractionator estimated the total number of RGCs by relating the number counted in the random counting frames to the sectional volume and then multiplying it by the reference volume.

## Acknowledgements

Grant support was provided by the National Eye Institute (R01 EY024984 to JSM), Indiana Department of Health Brain and Spinal Cord Injury Fund (JSM), an IU Collaborative Research Grant from the Office of the Vice President for Research (JSM), an award from the IU Signature Center for Brain and Spinal Cord Injury (JSM), an IUPUI Graduate Office First Year University Fellowship (KBL), and the Purdue Research Foundation Fellowship (KBL). This publication was also made possible with partial support from Grant # UL1TR002529 (A. Shekhar, PI) from the National Institutes of Health, National Center for Advancing Translational Sciences, Clinical and Translational Sciences Award (KB). Additional grant support was provided by IACUC #7611and Department of Defense grant #W81XWH-12-1-0617 (SW).

